# Yeasts prefer daycares and molds prefer private homes

**DOI:** 10.1101/2024.11.20.624568

**Authors:** Håvard Kauserud, Pedro M. Martin-Sanchez, Eva Lena Estensmo, Synnøve Botnen, Luis Morgado, Sundy Maurice, Klaus Høiland, Inger Skrede

## Abstract

Indoor fungi can cause negative health effects due to the production of toxins or volatiles that trigger the immune system of the occupants. To what degree indoor fungi (mycobiomes) differ between buildings with different usage is poorly known. Here, we compare the indoor mycobiomes in 123 children’s daycare centers and 214 private homes throughout Norway, as revealed by metabarcoding of DNA extracted from dust samples collected by community scientists. Although the fungal richness *per se* was similar in dust samples from daycares and homes, the fungal community composition differed. Yeast fungi, distributed mainly across the orders Saccharomycetales, Filobasidiales and Tremellales, were proportionally more abundant in the daycares, while filamentous fungi, including spore-producing molds such as *Aspergillus, Penicillum* and *Cladosporium*, were relatively more abundant in homes. Number of occupants, which is considerably higher in daycares, correlated significantly with the fungal community shift. We hypothesize that the density of occupants and their age distribution drive the systematic difference of yeasts and filamentous fungi in the two building types.

**Importance:** Worldwide, people spend most of their time indoors; in their homes, workplaces, schools and daycares. The indoor environment can thus largely influence our health. In this study, we revealed that number of occupants and possibly their age distribution are important explanatory variables for the fungal communities of the indoor environments in homes and daycares. Knowledge of these communities are important for further studies on how fungal diversity affect our health.

## Introduction

Within buildings, conditions for microbial growth are generally harsh due to limited humidity and scarce nutrient availability. However, some microorganisms are adapted to these adverse conditions and can grow and proliferate indoors. Molds and yeasts, both polyphyletic assemblages representing different fungal growth forms, are especially tolerant for the harsh indoor conditions and are often found in surveys of indoor fungal communities [1-9]. Molds are known to affect our health through the volatiles they produce or through aerially spread spores that may trigger our immune system or cause respiratory disease [10-12]. Many yeasts, such as *Candida* and *Malassezia*, are associated with the human body, where they mainly grow as commensals [13]. However, both yeasts and molds can cause superficial infections such as dandruff, atopic dermatitis/eczema, ringworm and nail infections [14], as well as serious infections in immuno-compromised people, e.g. invasive aspergillosis, mucormycosis and candidemia [15, 16]. The latter ones increased considerably during the COVID-19 pandemic [17, 18].

In addition to the fungi that can grow and survive indoor, fungal spores are transported indoors from outdoor sources and are detected in DNA-based surveys from the built environment [6, 7, 19-21]. Fungal spores spread easily by air into buildings through windows, doors and the ventilation system. Further, people and pets may function as vectors and transport fungal spores. The proportion of outdoor fungi spreading into buildings varies throughout the year, with a higher influx during the plant growth seasons, when fungi also are sporulating outdoors [5, 6, 20, 22].

In parts of the world, children of age 1-6 years spend considerable time inside daycare centers. Daycares are often characterized by a high density of people, which potentially influences air quality and humidity. High occupancy and low air quality have been suggested to allow higher yeast diversity in a study where yeasts were cultured from schools in Poland [3]. In Norway, outdoors play is highly evaluated and children in daycares spend up to 70% and 31% of their time outside during the summer and winter, respectively [23, 24]. Thus, outdoor materials, such as sand, soil, dust and plant debris, might easily be brought into daycares, constituting important biomass inputs for the indoor environment. Other elements usually not present in daycares, like potted plants and pets, are more common in homes, where the number of occupants is generally lower. In these respects, daycares may represent somewhat different environmental conditions for indoor fungal growth than homes. The indoor mycobiomes of daycares and private homes in Norway have previously been surveyed in separate studies, revealing a high prevalence of molds and yeasts in both building types [7, 8]. However, a direct comparison between these two settings is still lacking.

The main differences between the homes and daycares are the number of occupants and their age distribution, while the buildings themselves often can be similar, including similar architecture and same building materials. Logistically, it is challenging to obtain samples from a high number of buildings representative of a wide geographic region. In this study we therefore used a community science approach, recruiting inhabitants or daycare personnel to collect dust samples in a predefined simple manner, which allowed us to obtain a high number of samples throughout Norway for statistical comparisons. The key objectives of this study was to compare indoor dust mycobiomes from homes and daycares distributed throughout Norway. We aimed (i) to reveal whether different indoor mycobiomes can be found in the two building types and which fungal groups may differ, as well as (ii) to identify the factors that may be associated with these differences.

## Results and discussion

We compared two DNA metabarcoding datasets of indoors and outdoors dust samples from 214 homes and 123 daycares from throughout Norway (Appendix Figure A1), all collected in the spring 2018 [7, 8]. A weak, but significant difference in indoor fungal richness between the two building types was detected; we obtained on average 160 and 149 OTUs per sample for the indoor samples from homes and daycares, respectively (*t*-test, *p* = 0.02; Fig. 1a). Further, for homes, the fungal richness within the buildings was significantly higher than in the dust samples collected outdoors (*p* = 1.4e-14). Comparably, this increase was not significant for daycares (*p* = 0.34; Fig. 1a). In total, the daycare dataset had more OTUs than the homes dataset (7,419 and 6,408 OTUs, respectively; Fig. 1b). For both homes and daycares, only 11-12% of the fungal OTUs appeared uniquely outdoors, while 41-47% were uniquely found indoors. Other dust-mycobiome studies have also observed a higher diversity of fungi indoors, including the research conducted by Barberán et al. (2015) [25] and Yamamoto et al. (2015) [26]. This phenomenon can be explained by the fact that many outdoor fungi have the ability to enter buildings, while the reverse is apparently not the case to the same degree (Fig. 1b). Hence, the outdoor environment represents a major source of inoculum to the indoor environment, as also observed in previous studies [21, 25, 27]. In addition, the 49% of indoor fungi (OTUs) were found in both types of buildings, while 20% and 31% of them were uniquely associated with homes and daycares, respectively (Appendix Figure A2).

**Figure 1.**
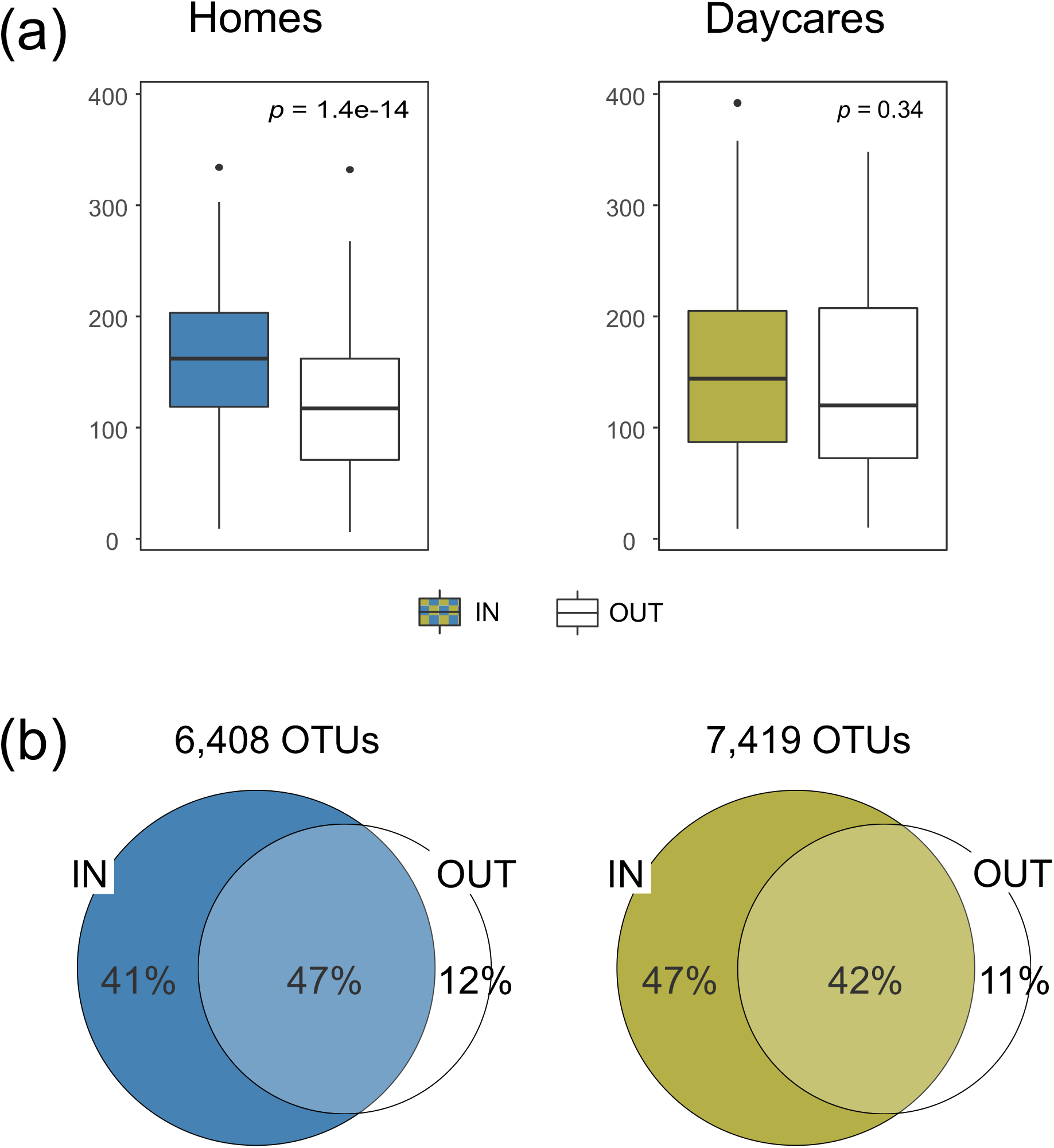
Comparison indoor vs. outdoor samples for each type of building: a) Richness per sample, and b) Number of OTUs and overlaps. All statistics were calculated from a rarefied matrix that includes 9,107 OTUs and 1,169 dust samples collected from homes (*n* = 636) and daycares (*n* = 533). Significance of richness differences between outdoor and indoor samples was assessed by *t*-test.

The community composition of the indoor mycobiomes was distinctly different in daycares and homes (Fig. 2a). The building type (daycare vs. homes) accounted for most of the variation in the indoor mycobiomes (6.3%), followed by the number of occupants (4.2%), being generally higher in daycares. Hence, our results suggest that the number of occupants, and possibly their age profiles, are important drivers for the differences in indoor dust mycobiomes between daycares and homes. Previous research has also reported higher airborne fungal loads (measured in colony forming units per m^3^) in daycares compared to homes [32]. In addition, the ventilation system of the building (balanced versus mechanical or natural) was of some significance (Fig. 2b).

**Figure 2.**
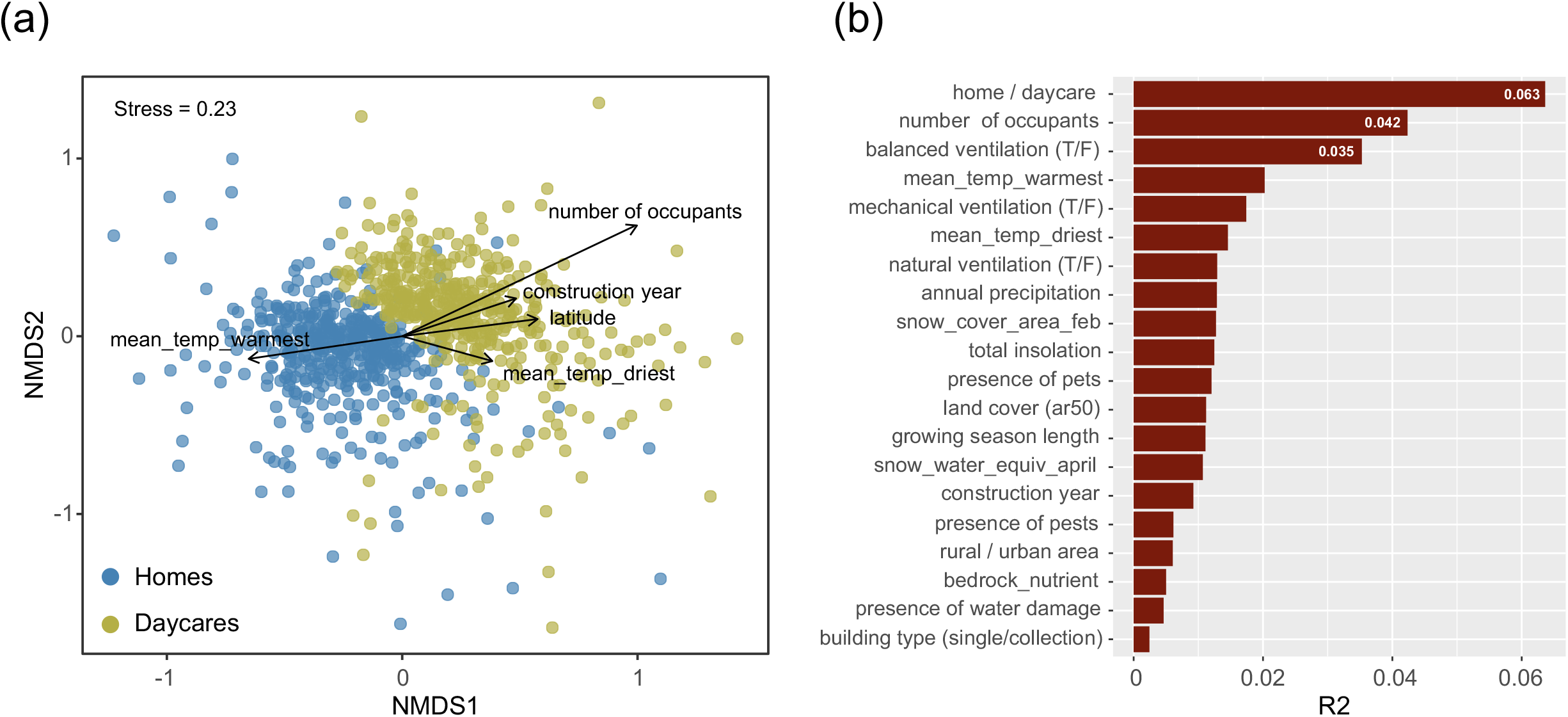
Factors correlated with compositional changes the indoor mycobiome in homes and daycares. a) NMDS ordination plot displaying compositional variation between homes and daycare centers. Each point represents one dust sample, and its color indicates its origin (home vs. daycare). b) The main explanatory variables for the observed variance according to PERMANOVA results (R2 values with *p* < 0.001). Statistics were based on the indoor rarefied matrix that includes 8,181 OTUs and 839 dust samples collected from homes (*n* = 428) and daycare centers (*n* = 411).

A high number of other factors were significantly correlated to the mycobiome composition, but accounted only for small proportions of the variation (Fig. 2b). For example, climate variables related to outdoor temperature and precipitation each explained less than 2.1% of the variation in the indoor mycobiome composition. The fact that the included environmental factors only account for a small part of the variation in community composition is a common feature in fungal community studies. The assembly process of fungal communities is probably strongly influenced by random processes, such as spore dispersal and colonization [28, 29], making exact predictions of mycobiome composition difficult. Furthermore, there is a high temporal (within-year) variation in fruiting and sporulation of outdoor fungi, especially in temperate regions, which is also reflected in the indoor mycobiomes due to the influx of spores [6, 30, 31]. In our previous temporal study of the mycobiomes in two daycares [6], dust samples were collected throughout a year in order to evaluate the effect of seasonality on the indoor mycobiomes using DNA metabarcoding. In this study, a strong seasonal pattern was observed in the mycobiome composition, with higher fungal richness in summer and fall. Hence, in analyses of indoor fungi, it is important to consider the temporal variability by obtaining samples at approximately the same time or conducting repeated sampling. In our study, the samples were collected throughout Norway at the same time period (April-May). Thus, even if the climate varies across the country, both the daycare and the home dataset are affected by the same climate variables.

We observed distinct differences in the taxonomic composition between the two building types (Fig. 3a). The orders Saccharomycetales, Filobasidiales and Tremellales, were proportionally more abundant in daycares. Further, on genus level, ascomycetous yeasts, like *Saccharomyces, Candida* and *Debaryomyces*, as well as basidiomycetous yeasts like *Cryptococcus, Filobasidium, Malassezia, Naganishia* and *Rhodotorula*, were proportionally more abundant in daycares compared to homes (Fig. 4, *t*-test *p* < 10E-5). In homes, saprotrophic and plant pathogenic filamentous ascomycetes in the orders Capnodiales, Dothideales, Eurotiales and Helotiales, were relatively more abundant (Fig. 3a). These orders include mold genera such as *Alternaria, Aspergillus, Cladosporium* and *Penicillium*, all proportionally more abundant in homes (Fig. 4). In contrast, the two mold genera *Wallemia* (Basidiomycota) and *Mucor* (Mucoromycota) were proportionally more abundant in daycares (Fig. 4). *Wallemia* includes many food contaminants [33] whereas *Mucor* includes soil saprotrophs that are commonly associated with food production and food spoilage [34]. Indicator species analysis also supported these findings, and identified some yeasts (*Filobasidium, Cryptococcus*, Saccharomyces and *Cyberlindnera*) and Mucor species as the strongest daycare indicators (IndVal > 50%), and the typical molds (*Penicillium, Alternaria, Aspergillus, Cladosporium* species) as home indicators (Appendix Table A2).

**Figure 3.**
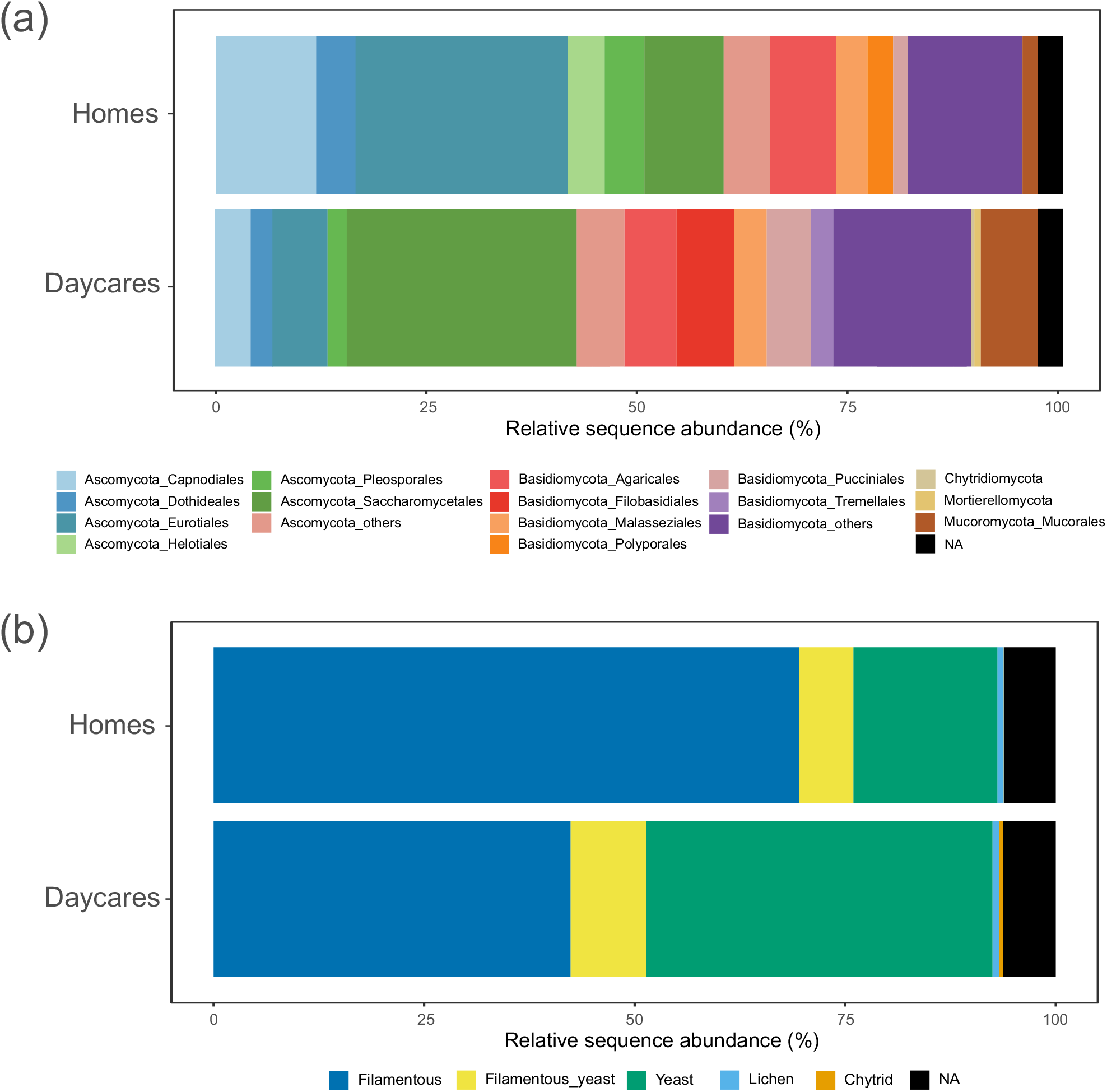
Relative sequence abundance of fungi detected in indoor dust samples from homes and daycares. a) The most abundant Orders sorted by Phyla. The less abundant Orders were collapsed and labelled as “Phylum_others”. b) Annotation of the main fungal growth forms: filamentous, filamentous and yeast (dimorphic fungi), yeast, lichen, and chytrid. NA: not assigned.

**Figure 4.**
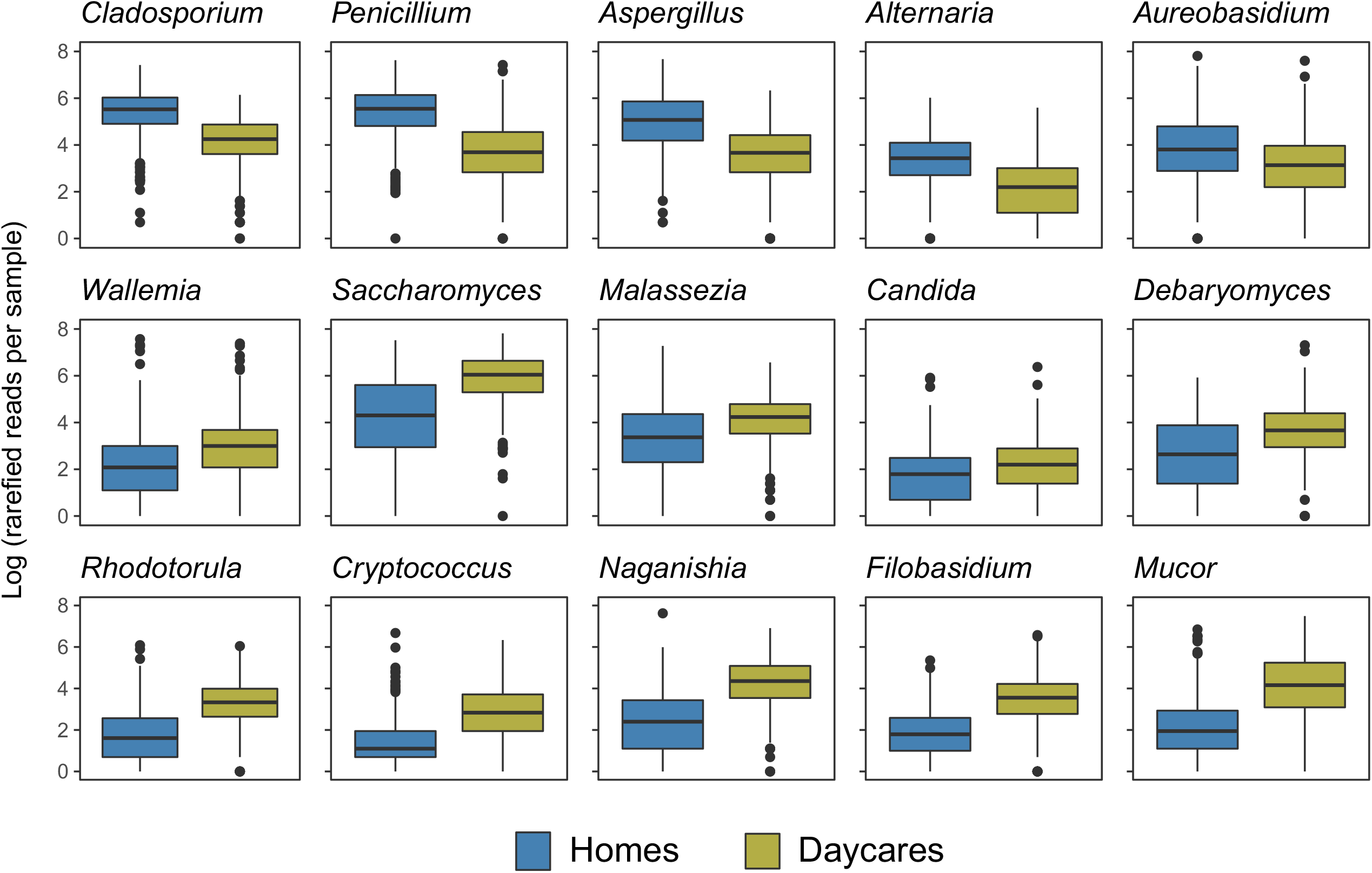
Selected genera showing significant differences (*p* < 10E-5; *t*-test) in abundance (Log (rarefied reads per sample)) when comparing indoor samples from homes (*n* = 428) and daycares (*n* = 411).

When annotating the OTUs in the final rarefied matrix (6,971 of 9,107 OTUs; 76.5%) into growth forms, we observed a clear difference in the distribution of yeasts, mycelial fungi and dimorphic fungi between the two building types (Fig. 3b), where yeast are relatively more abundant in daycares while mycelial fungi are relatively more abundant in homes. Previous research has shown that indoor environments, such as healthcare centers [2], homes [35] and schools [3, 36], exhibit high yeast diversity. While Marques do Nascimento et al. [2], Hashimoto et al. [35] and Ejdys et al. [3] specifically investigated yeasts by culturing, Park et al. [36] conducted metagenomic sequencing of all organisms in 500 classrooms. Both approaches identified a substantial level of yeast diversity including the genera *Candida, Debaryomyces, Rhodotorula, Cryptococcus, Naganishia, Filobasidium* and *Cyberlindnera*.

We suggest two different hypotheses that may explain the observed proportional difference between molds and yeasts in the two building types. First, more yeasts may be associated with young children, driving the difference. It has been documented that children have a more diverse fungal skin community compared to adults [37], including genera like *Aspergillus, Epicoccum, Cladosporium, Candida, Rhodotorula, Cryptococcus* and *Phoma*, in addition to the obligatory lipophilic yeast genus *Malassezia* that dominates on the skin of adults [37]. Moreover, the higher density of people *per se* may drive the proportional difference, since yeasts are more associated with the human body than molds [38]. Moreover, Adams et al. [9] reported a significant overlap between the mycobiomes associated with indoor environmental samples (dust and surfaces) and those from the occupants’ skin. Several fungal genera with yeast growth can also be found in the gastrointestinal tract, including yeast fungi such as *Candida, Malassezia* and *Saccharomyces* [39, 40]. A higher density of people may therefore lead to a proportional difference between yeasts and molds, which may be mediated in part by the deposition of occupants’ skin dead cells on the indoor surfaces. There seemed to be an even stronger difference in community composition between homes and daycares with many children (Fig. 2a), which may further support the latter hypothesis. However, to be able to conclude on this topic, more in-depth studies with cross factorial, balanced study design, tentatively also including investigations of the skin/body mycobiome, are needed.

Overall, there was a striking difference in the relative distribution of yeasts and filamentous fungi, where yeasts were proportionally more abundant in daycares and vice versa. Whether this difference is directly coupled to health effects is unknown. Molds have been shown to cause asthma and other respiratory diseases in humans in moist environments [41, 42]. Furthermore, moisture in homes, in addition to the level of fungal spores outdoors, were the best predictors to indoor fungal spore concentrations in 190 homes in Paris, France [43]. Moisture in schools, but not microbes, was the best predictor of respiratory problems in school children in the Netherlands and Finland [44]. However, a recent birth cohort study in Finnish homes reported that early-life exposure to home dust mycobiomes do not have clear negative or positive effects on asthma development in children [45]. Despite the clear association between some yeasts (e.g. *Malassezia* and *Candida*) and skin disorders (atopic dermatitis and mucocutaneous candidiasis, respectively) [14], some studies have pointed out a potential protective role of the dust yeast exposure against allergies and asthma in children [46]. Thus, the marked difference in the proportional abundance of molds and yeasts in the different building types may not lead to negative effects for the occupants. To gain further insight on this topic, future studies should assess inhabitant’s health status coupled to the indoor mycobiomes.

## Materials and methods

### Context and original datasets

We compare two datasets that previously have been analyzed and published separately [7, 8]. Since the sampling scheme, material and methods were thoroughly described in the original publications, we provide a condensed version here. Altogether 271 homes and 125 daycares throughout Norway were originally selected for sampling. However, the combined dataset of this study includes 428 indoor samples from 214 homes and 411 indoor samples from 123 daycares to include a more balanced numbers of indoor samples and correct the overrepresentation of Oslo area in the original home dataset. During spring 2018, inhabitants (homes) or personnel (daycares) collected dust samples on doorframes at three specific locations: (1) the main entrance outdoors, (2) main central room (living room in homes), and (3) bathroom. Large daycares sampled from two main central rooms and two bathrooms. The dust samples were obtained using the same sampling kits including sterile FLOQSwabs (Copan Italia spa, Brescia, Italy) and instructions. The returned swabs were stored at -80 ºC until DNA extraction. The inhabitants/personnel also provided metadata about the buildings such as number of occupants, building features and previously reported pests and water damages by responding to a questionnaire. In addition, based on the geographical coordinates of the buildings, data for some relevant environmental variables related to climate, geology and topography were extracted from WorldClim 2 or provided by [47] (see Appendix Table A1 for metadata). In brief, the DNA metabarcoding workflow included five steps: (i) DNA extraction from the swabs using chloroform and the EZNA Soil DNA Kit (Omega Bio-tek, Norcross, GA, USA); (ii) PCR amplification of the ITS2 region using the primers gITS7 [48] and ITS4 [49], both including sample specific tags at the 5’-end; (iii) clean up and normalization of PCR products using SequalPrep Normalization Plates (Thermo Fisher Scientific, Waltham, MA, USA), and subsequent pooling of 96 uniquely barcoded samples including technical replicates, negative samples (unused swabs), extraction blanks, PCR negatives, and a mock community; and (v) library preparation and 250 bp paired-end MiSeq Illumina sequencing carried out at Fasteris SA (Plan-les-Ouates, Switzerland).

### Bioinformatics

The bioinformatic analyses for the combined dataset from homes and daycares, whose raw sequences are available on ENA at EMBL-EBI (https://www.ebi.ac.uk/ena/browser/view/PRJEB42161) and Dryad (https://doi.org/10.5061/dryad.sn02v6×5s), respectively, were performed as described in Martin-Sanchez et al. (2021) and Estensmo et al. (2022) with slight modifications. Shortly, raw sequences were demultiplexed using CUTADAPT [50] and sequences shorter than 100 bp discarded. DADA2 [51] was used to filter low quality reads, error correction, merging in contigs and chimera removal. ITSx [52] was used to exclude the non-fungal sequences and trim the conserved regions of flanking rRNA genes. To account for intraspecific variability [53], the generated amplicon sequence variants (ASVs) were clustered into operational taxonomic units (OTUs) using VSEARCH [54] at 97% similarity. LULU [55] was used with default settings to correct for potential OTU over-splitting. Taxonomy of OTUs was assigned using the BLASTm algorithm [56] against the UNITE and INSD dataset for fungi (Version 04.02.2020) [57] Ecological trophic modes and guilds for the identified taxa were annotated using the FUNGuild tool [58]. OTUs with less than 10 reads and those that were not assigned to the kingdom Fungi were discarded from downstream analyses. For comparing daycares and homes, we downscaled the original datasets by excluding 2 daycares and 57 homes, hereby providing a more balanced dataset in terms of geographical location (15 homes per municipality maximum), collection date (all samples in April-May 2018) and number of indoor samples from homes vs. daycares (428 vs. 411 in the rarefied matrix). The OTU table was rarefied to 2,540 reads per sample using the function *rrarefy* of the VEGAN R package [59], keeping the majority of samples (only 18 samples were excluded). The final quality-filtered and rarefied matrix, without technical replicates, negative controls and mock samples, contained 9,107 OTUs from 1,169 samples. Those OTUs with taxonomic assignment at species, genus or family level were further annotated into growth forms (filamentous, yeast, dimorphic, lichen and chytrid) based on literature surveys.

### Statistics

Initially, we assessed species richness (number of observed OTUs) per sample, as well as the total number of OTUs and their overlaps for the two types of building (homes vs. daycares) and compartments (indoor vs. outdoor). For comparison of the indoor mycobiomes, beta diversity was assessed with NMDS ordination of dust samples using *metaMDS*, Bray-Curtis dissimilarity index and 200 random starts in search of stable solution on the Hellinger-transformed rarefied OTU tables. Continuous environmental variables were regressed against NMDS ordination and added as vectors on the ordination plots using *gg_envfit* of the R package *ggordiplot* v 0.3.0 [60] to visualize their association with the indoor dust mycobiomes. To evaluate the correlation between environmental variables and the observed variance in fungal community composition, permutational multivariate analysis of variance (PERMANOVA; 999 permutations) was performed individually on each variable using *adonis2*. Relative abundances of taxa at order and genus level were assessed to highlight the differences between homes and daycares. To reveal significant associations (*p* < 0.05) between OTUs and the type of building, an indicator species analysis was performed using *multipatt* of the R package *indicspecies* [61]. Significant differences in the variance of species richness per sample and the relative abundances of selected genera were evaluated with the analysis of variance (ANOVA) and *t*-test.

### Data availability

Our initial combined OTU table, as well as the final rarefied matrix for fungi, are available at Zenodo (https://doi.org/10.5281/zenodo.14049800) together with information about metadata of environmental variables, taxonomic assignment, as well as annotations of trophic modes/guilds and growth forms.

## Acknowledgements

We acknowledge the personnel of the daycare centers and the inhabitants of the homes for sampling and for providing metadata of the building and occupancy features. Mycoteam AS contributed to the sampling and provided sampling equipment. The research was financially supported by the University of Oslo, the Norwegian Asthma and Allergy Association (NAAF), and the European Union’s Horizon 2020 research and innovation program (Marie Skłodowska-Curie Individual Fellowship to PMM-S; grant agreement *MycoIndoor* No 741332). PMM-S also thanks the grant PID2021-123184OA-I00 funded by MCIN/AEI/ 10.13039/501100011033 and ERDF - A way of making Europe.

## CRediT authorship contribution statement

Håvard Kauserud: Conceptualization, Writing – original draft, Writing – review & editing, Supervision, Funding acquisition. Pedro M. Martin-Sanchez: Conceptualization, Investigation, Methodology, Formal analysis, Data curation, Visualization, Writing – review & editing, Funding acquisition. Eva Lena Estensmo: Conceptualization, Investigation, Methodology, Writing – review & editing, Funding acquisition. Synnøve Botnen: Writing – review & editing. Luis Morgado: Software, Formal analysis, Writing – review & editing. Sundy Maurice: Methodology, Resources, Writing – review & editing. Klaus Høiland: Formal analysis, Writing – review & editing. Inger Skrede: Conceptualization, Writing – review & editing, Supervision, Funding acquisition.

## Appendix (supplementary material)

**Figure A1**. Location and distribution of the private homes (214) and daycares (123) selected throughout Norway for this study.

**Figure A2**. Overlap of indoor dust mycobiomes (8,181 OTUs) between homes and daycares.

**Table A1**. Variables analyzed in this study

**Table A2**. OTUs identified as indoor indicator fungal species (IndVal > 50%; *p* < 0.05) for homes and daycares using the R package *indicspecies*.

